# Biological Control Potential of Ectomycorrhizal Fungi against *Fusarium circinatum* on *Pinus patula* seedlings

**DOI:** 10.1101/652511

**Authors:** Veronique Chartier-FitzGerald, Joanna Felicity Dames, Greer Hawley

## Abstract

The South Africa forestry industry, covering 1.3 million hectares, is dependent on exotic pine and eucalyptus species. Nursery seedlings are not inoculated with ectomycorrhizal (ECM) fungi or other beneficial microbes. *Fusarium circinatum* is an economically important pathogen affecting seedling survival. The purpose of this investigation was to assess to determine the effects of ectomycorrhizal fungal inoculation on *Pinus paulta* seedling growth and resistance to the fungal pathogen *F. circiantum*. Explants from ECM basidiocarps, collected from *Pinus* stands, were plated onto MMN medium to obtain isolates which were verified using molecular techniques. These isolates were identified as *Boletus edulis f. reticulatus, Lactarius quieticolor, Suillus granulatus* and an additional *Suillus* isolate. *P. patula* growth in the presence of the pathogen *F. circinatum* was significantly increased and promoted by the *L. quieticolor* and *Suillus* isolates. Inoculation of seedlings in the nursery would ensure the production of stronger healthy plants which may be more tolerant to fusarial infection, increasing survival in the plantation.

## Introduction

Since the first detection of *Fusarium circinatum* (teleomorph= *Gibberella circinata*) in South Africa on *Pinus patula* seedlings in 1990 ^(1)^ this pathogen has become a major problem in production nurseries. It causes damping off, root and collar rot, and tip dieback, often resulting in large scale seedling mortality ^(2)^. Due to these significant losses in yield and productivity this pathogen was characterised as the largest limitation to commercial forestry ^(1;2;3)^. In South Africa, the most susceptible of the *Pinus* species is *P. patula* ^(4)^ but it is also the most commonly planted species as more resistant species such as *P. radiata* and *P. elliottii* produce wood of a poorer quality ^(3; 5)^.

The impact of *F. circinatum* is not only felt within the nurseries but also in plantations, where significant losses are experienced after outplanting. Crous ^(6)^ determined that approximately 42% of all recently planted seedlings over 16 sites died due to infection with *F. circinatum*. The majority of the mortalities occurred within the first 30-140 days after planting, with a mean seedling survival of 36 – 53% after 1 year. This implies that at least half of all field seedling death is due to infection. The *F. circinatum* related mortalities of *P. patula* result in losses in excess of R11 million per year to the industry, and in excess of R12 million a year when *P. radiata*, the second most susceptible *Pinus* species, is included ^(3;4)^.

The high levels of *F. circinatum* occurrence in South African nurseries are attributed to contaminated nursery containers, irrigation and in some cases growing media and plants ^(3;7)^. Infection rates have also been recorded to rise with an increase in the amount of nitrogen (N) given to seedlings ^(3)^. The current approaches for control of *F. circinatum* include improved nursery sanitation and long-term strategies such as cloning, hybridisation breeding and selection to produce hybrids and trees of increased resistance to the pathogen. While the increased nursery sanitation strategy is showing promise ^(7)^ there are a number of downsides to the cloning and hybrid programs. There is therefore a need for a more economical and less time-consuming alternative for the control of this pathogen. One solution may lie in the early inoculation and establishment of ectomycorrhizal (ECM) fungi.

Most ECM fungi will inhibit non-mycorrhizal fungal growth via indirect means or via direct competitive exclusion for both space and nutrients. The presence of ECM fungi decreases the rate of pathogenic infection as a barrier (the ECM mantle) along the root tip is formed, preventing pathogenic root infections ^(8;9)^. Non-mycorrhizal roots are more prone to infection ^(10;11)^, especially succulent root growth which is predisposed to infection from *F. circinatum* ^(3)^. Competition for nutrients has been demonstrated between non-pathogenic strains of *Fusarium* against their pathogenic counterparts. It is highly probable that the ECM fungi would similarly compete for nutrients and therefore act as effective biological control agents of root pathogens ^(12)^.

More direct inhibition methods involve the production of antagonistic antifungal compounds such as chitinases or phenols ^(11;13;14)^. ECM fungi have also been found to be effective against different strains of bacterial wilt in nurseries, decreasing disease rates from 72 to 39% ^(15)^. Suppressive effects such as these are considered to be important to the survival of *Pinus* seedlings in nursery and forestry environments (Smith and Read, 2008).

The aims of this study are to select, culture and identify local ECM fungi, and determine whether ECM fungal inoculum can increase *P. patula* seedling growth when challenged with the pathogen *F. circinatum*.

## Methods

### Isolation of ECM fungi from selected fruiting bodies

ECM fruiting bodies were collected from *Pinus* stands in the highlands of Grahamstown in April 2015. The collected fruiting bodies were visually identified using field guides ^(16;17)^. Explants from within the fertile cap of the fruiting bodies were extracted using sterile technique and placed onto Modified Melin-Norkrans (MMN) ^(18)^ agar and incubated at 28°C.

### Molecular Identification

From the sporocarp material collected above, molecular identifications were performed to confirm the visual identifications. This was achieved using the ZR Fungal/Bacterial DNA MiniPrep kit. The successful extraction of DNA was confirmed by visualisation on a 1% agarose gel stained with ethidium bromide and photographed using a UV Transilluminator.

The Internal Transcribed Spacer (ITS) region of rDNA gene was amplified via PCR using the primers ITS1F and ITS4 ^(19)^. A 25 µl reaction contained: 12.3 µl of sterile water, 5 µl of 5X Kapa Hifi GC Buffer, which contained 2mM MgCl_2_ (1X), 5 µl of template DNA, 0.75 µl of 10 mM dNTP mix, 0.75 µl of each primer and 0.5 µl of the Kapa Hifi HotStart DNA polymerase. PCR amplification was conducted under the following conditions: 95ºC for 5 mins, followed by 25 cycles of denaturation at 98ºC for 30 s, annealing at 47ºC for 45 s and extension at 72ºC for 1 min followed by a final extension at 72ºC for 7 mins. Sequencing was performed at Inqaba Biotechnology Industries (Pty) Ltd. Pretoria. The resulting Sanger sequences were analysed using Mega 7 ^(20)^ and submitted to the GenBank (www.ncbi.nlm.nih.gov/BLAST/) ^(21)^ and UNITE ^(22)^ databases once identified to genus level using BLAST.

#### Fusarium circinatum isolates

Isolates of *F. circinatum* were provided by the Forestry and Agricultural Biotechnology Institute (FABI), University of Pretoria, South Africa. The 5 isolates (CMWF 666 (VCO 21), CMWF 594 (VC08), CMWF 623 (VCO25), CMWF 701 (VCO30), CMWF 621 (VCO6)) provided, were originally collected from local South African *Pinus radiata* nurseries ^(2)^. These isolates were cultured on Potato Dextrose Agar (PDA) at 28°C. For brevity the isolates will be abbreviated to FC 666, FC 594, FC 623, FC 701, FC 621.

### Antifungal activity assay

To visualise the interaction between the ECM fungal isolates and the *F. circinatum* isolates an antifungal activity dual assay was conducted. Plates were divided into 4 sections and inoculated with plugs of the different *F. circinatum* isolates, above the central intersecting lines. The remaining 4 quadrants were inoculated with one of the ECM fungal isolates. Two replicates per ECM fungal isolate per *F. circinatum* strain were used. A visual assessment of any growth inhibition was made.

#### Greenhouse trial

##### Pinus patula seedlings

The seedlings were grown from *P. patula* seeds provided by the Institute for Commercial Forestry Research (ICFR), Pietermaritzburg, South Africa. Seeds were surface sterilised in 2% sodium hypochlorite (commercial bleach) and rinsed in sterile water. They were grown under controlled conditions in a 1:1 mix of sterile perlite and vermiculite for a period of 2 months before use in the trial.

##### Inoculum preparation

The different ECM isolates were grown in 250 ml of MMN broth at 25ºC for a month to allow for maximum growth. On the day of the pot trial the ECM fungal cultures were homogenized using an Ultraturex. To standardize the concentration of each ECM fungal inoculum the homogenate was measured spectrophotometrically at a wavelength 600 nm and adjusted with sterile water to match the lowest OD reading. The homogenized fungal mixtures were then added to 0.3% water agar to form 1 L of a gel inoculum each.

The *F. circinatum* isolates were grown on PDA for a minimum of 2 weeks before harvesting for spores. The spores were then added to 20 ml of sterile 15% glycerol and vortexed for 30 seconds to produce a homogenous solution. From this initial suspension a 1:100 dilution with sterile water was made; and spore concentrations were then determined using a Neubauer hemocytometer. For each conidial preparation made the viability of the spores was confirmed by spread plating 100 μl onto PDA plates. The percentage of spores which germinated was determined after 48 hours’ incubation at 25°C.

##### Greenhouse trial

Pots were sterilised with a 2% solution of sodium hypochlorite. The bottom of which were covered with surface sterilised pebbles to prevent soil loss and increase drainage. Each pot was filled half-way with a sterilised 2:1:1 mixture of compost, perlite and vermiculite. Seedling roots were placed in the appropriate ECM fungal inoculum mixture for approximately 10 minutes before planting, and an additional 8 ml of this same mixture was added to the roots of each seedling once planted to ensure colonisation. The seedlings of the negative control were soaked and inoculated with sterile water agar mix.

One week into the trial, 1 ml *F. circinatum* spores at a concentration of 1 × 10^6^ ml^−1^ were added to the soil above the roots. The number of replicates and treatment configurations are illustrated in table 1. The initial heights were measured upon planting and subsequently measured and recorded weekly for 9 weeks. Seedlings were placed in a mycorrhizal research tunnel having an average temperature range of 25 to 35°C, pots were irrigated daily with UV treated water and grown under natural lighting.

**Table 1.** Summary of treatments and replicates for biological control trial

|  |  | Replicas per treatment |  |  |  |  |
| --- | --- | --- | --- | --- | --- | --- |
| Ectomycorrhizal |  | *Fusarium circinatum* strains |  |  |  |  |
| Isolates | Control | FC 594 | FC 621 | FC 623 | FC 666 | FC 701 |
| Negative Control | 10 | - | - | - | - | - |
| Positive Controls | - | 10 | 10 | 10 | 10 | 10 |
| Lactarius | 10 | 10 | 10 | 10 | 10 | 10 |
| Boletus | 10 | 10 | 10 | 10 | 10 | 10 |
| Salmon Suillus | 10 | 10 | 10 | 10 | 10 | 10 |
| Suillus | 10 | 10 | 10 | 10 | 10 | 10 |

After 9 weeks the seedlings were carefully removed from their containers and transported to the laboratory for further analysis. The roots were severed from the shoots and wet weights recorded.

The percentage colonisation for each seedling was determined using a modified line intersect method ^(23)^. Every time a colonised root came into contact with a gridline this was recorded along with its mark representing a gridline intersection. The percentage colonisation was the percentage of intersection marks which represented roots with ECM fungal colonisation from the seedling’s overall number of intercepts. The roots were then dried in an oven at 60°C for 3-4 days and the resulting dry biomass was recorded.

#### Statistical analysis

A Shapiro test was performed on the percentage colonisation and root dry weight data to determine normality. The null hypothesis was rejected for both showing the data was non-parametric. Thus, a Kruskal-Wallis Analysis of Variance (ANOVA) was performed and a pairwise Wilcox test was performed to determine significant difference between treatments.

Seedling growth was analysed using a repeated measures ANOVA. A least-squares means pairwise comparison with a Tukey adjustment was performed to compare treatments to one another and determine significant differences. All statistical analyses were performed using RStudio Version 0.99.903 ^(24)^.

## Results

### Isolation and identification of ECM fungi from fruiting bodies

The fruiting body samples collected were field identified as *Boletus edulis, Suillus granulatus, Suillus salmonicolor* and *Lactarius deliciosus* (Table 2), these isolates will be referred to as Boletus, Suillus, Salmon Suillus and Lactarius for the remainder of the study. Salmon Suillus did not resolve to a satisfactory molecular identification. Unfortunately due to limited amounts of sample it was not possible to repeat the sample to resolve the specific species and provide a better sequencing result.

**Table 2:** Summary of BLAST results for the ECM sporocarps.

| Sporocarp | Accession<br>Number | Genera | Aligned<br>with | Percentage<br>Identity | E-value | Query<br>Coverage |
| --- | --- | --- | --- | --- | --- | --- |
| Boletus | MG806927 | *Boletus edulis f.*<br>*reticulatus* | KY595992 | 95 | 0.0 | 100% |
| Lactarius | MG833316 | *Lactarius quieticolor* | KX610696 | 100 | 0.0 | 100% |
| Salmon<br>Suillus | MG806929 | *Suillus* | KX170996 | 91 | 1e-155 | 100% |
| Suillus | MG806928 | *Suillus granulatus* | KU721244 | 99 | 0.0 | 100% |

### Fusarium circinatum antifungal activity assay

The *F. circinatum* growth was visibly inhibited and the pathogen actively avoided the ECM fungi most notably in the presence of Lactarius and Suillus. While the inhibition is not as dramatically visible, both Salmon Suillus and Boletus also resulted in growth avoidance of the pitch canker isolates.

### Greenhouse trial

The linear mixed effects model showed that the different treatments of ECM fungi and the combination of ECM fungi and *F. circinatum* had significant effects on the growth of the seedlings.

From figure 1 it is can be seen that inoculations with the fungus Lactarius produced the highest growth in comparison to the other ECM fungi and control treatments and more importantly continued to promote and increase the growth of seedlings inoculated with the different *F. circinatum* strains. It was the only ECM fungus to significantly increase the growth of the *P. patula* seedlings infected by the pitch canker strains; specifically, it was significant against the strain 594. This significant growth in the presence of *F. circinatum* is reflected in the antifungal assays, where Lactarius had the most visible inhibition of *F. circinatum* growth.

**Figure 1.**
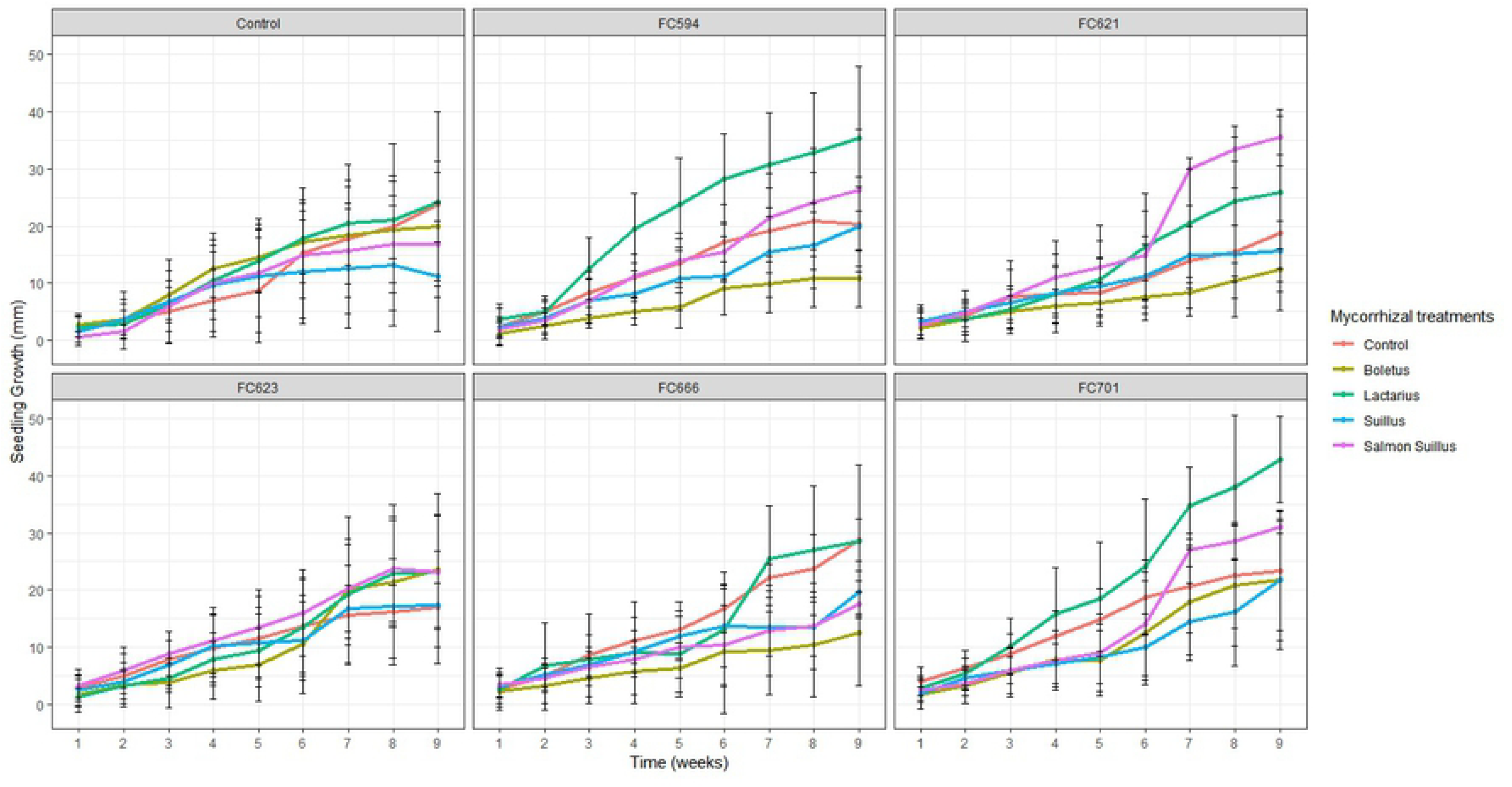
Greenhouse trial average weekly growth of the *P. patula* seedlings for each treatment over a period of 9 weeks, separated according to *Fusarium circinatum* treatment. Error bars represent ± standard error. ANOVA linear mixed effects model (Intercept: F-value = 932.1432, P-value <0.0001, ECM fungi: F-value = 12.1650, P-value <0.0001, *F. circinatum* F-value 3.0175, P-value = 0.0114, ECM fungi + *F. circinatum* F-value = 20.441, P-value = 0.0061)

For most treatments the application of Salmon Suillus produced the second highest average *P. patula* seedling growth (Figure 1). Since Salmon Suillus produced much less visible inhibition of *F. circinatum in vitro* it is highly likely that it increased seedling growth while inhibiting pathogenic infection using a different mechanism to Lactarius. Boletus and Suillus did not always increase the *P. patula* seedling’s growth in comparison to the *F. circinatum* controls. Although both ECM fungi did demonstrate visible *in vitro F. circinatum* inhibition, it is proposed that even though plant growth is not being increased, pathogen inhibition is occurring.

The higher levels of growth and colonisation when exposed to the pathogen *F. circinatum* indicate the benefits of using ECM fungal inoculum on *Pinus* seedlings. Some very low levels of colonisation did occur in the controls, although this is likely due to splashing from surrounding pots. While Lactarius had the highest seedling growth it had the lowest percentage colonisation, with significantly lower colonisation in the presence of some *F. circinatum* isolates, such as FC 666, 621 and 623 (Figure 2). On the other hand, Salmon Suillus had the highest percentage colonisation compared to other ECM fungi, especially with respect to treatments with FC 621 and FC 666. Boletus had the second highest levels of colonisation on average closely followed by Suillus.

**Figure 2.**
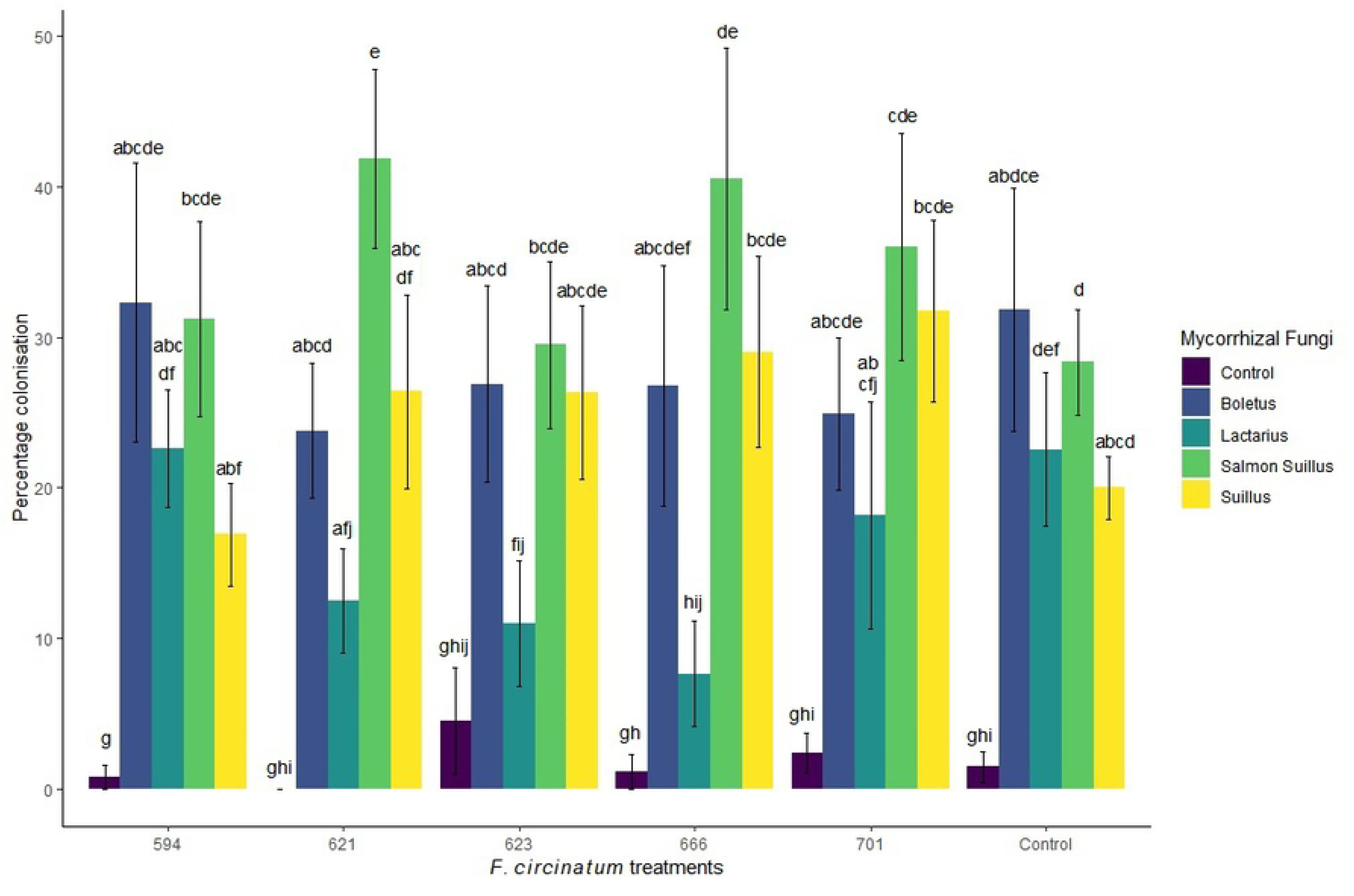
Average colonisation levels of *P. patula* seedlings after inoculation with ECM seedlings and exposure to *F. circinatum* (Kruskal-Wallis H (29, 270) = 130.02, p = 9.497e-15). Error bars represent ± standard error. Columns with the same letters are not significantly different from each other.

The colonisation levels for the most part did not correlate with root weight. Due to high variability, Suillus had the largest root biomass, which exceeded Lactarius and Salmon Suillus (Figure 3). Boletus had the second largest root biomass, as would be expected from its levels of colonisation. On the other hand Salmon Suillus had significantly smaller root biomass than either of these two treatments, on par with the *F. circinatum* control seedlings, while Lactarius gave variable results. Thus, while colonisation was high this did not correspond to higher root biomass.

**Figure 3.**
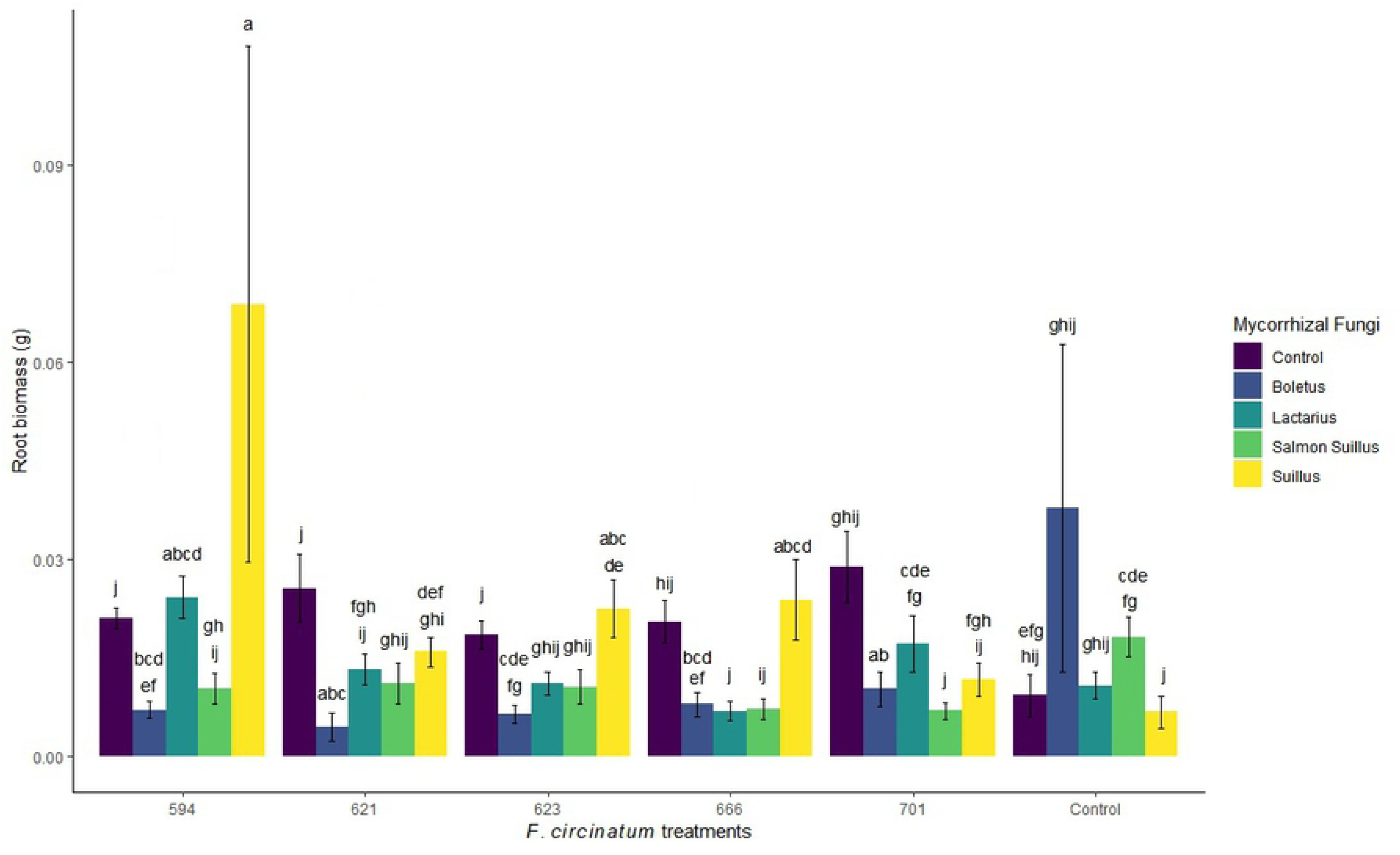
Average root biomass for each treatment (Kruskal-Wallis H (29, 270) = 117.38, p=1.36e-12). Error bars represent ± standard error. Columns with the same letters are not significantly different from one another

## Discussion

The ECM fungus Lactarius significantly improved the growth of the *P. patula* seedlings inoculated with pitch canker, especially isolate 594. While the increase in growth of plants subjected to other ECM + FC treatments were not significant, an increase in growth was still observed for all Lactarius and Salmon Suillus + FC strains treatments. Salmon Suillus had the highest growth and corresponding high levels of root colonisation. Thus any future work on aspects of ECM fungal biological control of *F. circinatum* would need to focus on both the Lactarius and Salmon Suillus isolates.

This increase in growth as a result of Salmon Suillus inoculation was unexpected based on the dual anti-pathogenic *in vitro* assay that was conducted, as Salmon Suillus showed little to no visible levels of inhibition against *F. circinatum.* This indicates that it employs an indirect mode of action through colonisation of roots, thereby limiting pathogen infection sites. ECM fungi form a mantle around each and every root tip they colonize which can be anything from 1-2 hyphal diameters thick to 30-40 diameters. Thus in order for a pathogen to enter the plant roots, it must now penetrate the ECM fungal mantle and the plant epidermal cell walls before infection can occur ^(8; 9)^. It is also well known that non-mycorrhizal roots, especially non-lignified growth, are targeted by *F. circinatum.* Thus, mycorrhizal colonisation is a viable form of biological control, particularly at the seedling stage. In a study performed by Branzanti ^(9)^ it was found that spores of the pathogens *Phytophthora cinnamomi* and *P. cambivora* were only detected on non-ECM colonised roots. In natural environments ECM fungi not only competitively exclude pathogens from infection sites but also compete for nutrients, such as litter patches. The ECM fungi use the carbon acquired from the host plant to rapidly colonise and grow within these nutrient sources and deplete the available nutrients in advance of other competitor organisms ^(24)^.

Conversely Lactarius, despite producing a significant increase in seedling growth, had the lowest level of colonisation. Thus, it is proposed that this ECM fungus exhibits a more direct form of inhibition against *F. circinatum.* This was demonstrated by the strong growth inhibition of *F. circinatum* observed in the *in vitro* dual assay. Similar results of low colonisation levels and inhibited *Fusarium* infection were observed by Mateos ^(25)^. Their study found that mycorrhizal formation was significantly decreased when *P. sylvestris* and *P. pinea* seedlings were co-inoculated with *Suillus luteus* and *F. oxysporium* or *F. verticillioides* yet fungal infection was still inhibited. This indicates that the inhibition of the pathogenic fungus was not related to competition for space but was rather due to a direct form of inhibition ^(25)^. Direct forms of inhibition include the production of antifungal enzymes like chitinases and ß-1,3-glucanases. Both of these are considered to be important enzymes in the lysis of fungal cell walls as chitin and ß-1,3-glucan are key fungal cell wall components and have been found in the cell walls of many plant pathogens ^(11;26)^. Mohan ^(11)^ found that while the amount of chitinase produced differed between ECM fungal species in dual culture *in vitro* experiments each species tested exhibited significant inhibition of different plant pathogens. ECM fungi also produce other anti-fungal compounds such as oxalic acids, phenolics, steroids and hydrogen peroxide ^(13;14;27;28)^. For example Duchnes ^(29:30)^ observed a six-fold reduction in the sporulation of the pathogen *Fusarium oxysporum* in the *Pinus resinosa’s* rhizosphere after inoculation with the ECM fungus *Paxillus involutus.* This was found to be due to ethanol-soluble compounds with fungal-toxic effects found in the rhizosphere 3 days after ECM fungal inoculation.

From the Suillus’ anti-fungal assay it was highly likely that it too inhibited *F. circinatum* infection via direct means even though the Suillus treatment did not increase the growth of the *P. patula* seedlings. Mateos ^(25)^ also reported cases of levels of ECM fungal colonisation, high fungal pathogen inhibition and absence of improved seedling growth.

It appears though that Boletus uses a more indirect form of pathogen inhibition, similar to Salmon Suillus, by aggressively colonising plant roots and increasing root biomass. It is also possible that the inhibition observed in the trial was due to increased levels of plant nutrition attained via ECM association. This would have allowed the host plant to disproportionately allocate resources to its defense mechanisms ^(31)^. Increased seedling growth seen as a result of ECM inoculation shows that overall the plant’s tolerance and resistance to infection from the presence of *F. circinatum* in the potting soil was decreased due to the biological control properties, direct or indirect, of the ECM fungi.

An interesting area of future study would be a combination of the two best performing ECM fungi, Salmon Suillus and Lactarius against the different *F. circinatum* strains.

In conclusion it can be seen from the trial conducted that of the successfully isolated ECM fungi, the two best isolates for inoculation of *P. patula* seedlings in South African nurseries would be *Lactarius quieticolor* and the *Suillus* isolate Salmon Suillus. Optimisation of fungal growth and development of an inoculum would be the next step in reaching this goal.

## References

1 Wingfield MJ, Hammerbacher A, Ganley RJ, Steenkamp ET, Gordon TR, Wingfield BD, Coutinho TA. Pitch canker caused by *Fusarium circinatum*—a growing threat to pine plantations and forests worldwide. Australasian Plant Pathology. 2008 Jul 1;37(4):319–34.

2 Steenkamp ET, Makhari OM, Coutinho TA, Wingfield BD, Wingfield MJ. Evidence for a new introduction of the pitch canker fungus *Fusarium circinatum* in South Africa. Plant Pathology. 2014 Jun;63(3):530–8.

3 Mitchell RG, Steenkamp ET, Coutinho TA, Wingfield MJ. The pitch canker fungus, *Fusarium circinatum*: implications for South African forestry. Southern Forests: a Journal of Forest Science. 2011 Jan 1;73(1):1–3.

4 Viljoen A, Wingfield MJ, Kemp GH, Marasas WF. Susceptibility of pines in South Africa to the pitch canker fungus *Fusarium subglutinans* f. sp. *pini*. Plant pathology. 1995 Oct;44(5):877–82.

5 Department of Agriculture, Forestry and Fisheries, Report on commercial timber resources and primary roundwood processing in South Africa (2014/15)

6 Crous JW. Post establishment survival of *Pinus patula* in Mpumalanga, one year after planting. Southern African Forestry Journal. 2005 Nov 1;205(1):3–11.

7 Van Wyk SJ, Boutigny AL, Coutinho TA, Viljoen A. Sanitation of a South African forestry nursery contaminated with *Fusarium circinatum* using hydrogen peroxide at specific oxidation reduction potentials. Plant disease. 2012 Jun;96(6):875–80.

8 Marx DH. Ectomycorrhizae as biological deterrents to pathogenic root infections. Annual review of phytopathology. 1972 Sep;10(1):429–54.

9 Branzanti MB, Rocca E, Pisi A. Effect of ectomycorrhizal fungi on chestnut ink disease. Mycorrhiza. 1999 Aug 1;9(2):103–9.

10 Ramachela K, Theron JM. Effect of ectomycorrhizal fungi in the protection of *Uapaca kirkiana* seedlings against root pathogens in Zimbabwe. Southern Forests. 2010 May 17;72(1):37–45.

11 Mohan V, Nivea R, Menon S. Evaluation of ectomycorrhizal fungi as potential bio-control agents against selected plant pathogenic fungi. JAIR. 2015 Feb;3(9):408–12.

12 Whipps JM. Microbial interactions and biocontrol in the rhizosphere. Journal of experimental Botany. 2001 Mar 1;52(suppl_1):487–511.

13 Suh HW, Crawford DL, Korus RA, Shetty K. Production of antifungal metabolites by the ectomycorrhizal fungus *Pisolithus tinctorius* strain SMF. Journal of industrial microbiology. 1991 Jul 1;8(1):29–35.

14 Yamaji K, Ishimoto H, Usui N, Mori S. Organic acids and water-soluble phenolics produced by *Paxillus* sp. 60/92 together show antifungal activity against *Pythium vexans* under acidic culture conditions. Mycorrhiza. 2005 Jan 1;15(1):17–23.

15 Gong M, Chen Y, Wang F, Chen Y. Inhibitory effect of ectomycorrhizal fungi on bacteria wilt of Eucalyptus. Forest Research. 1999;12(4):339–45.

16 Van Wyk SJ, Boutigny AL, Coutinho TA, Viljoen A. Sanitation of a South African forestry nursery contaminated with *Fusarium circinatum* using hydrogen peroxide at specific oxidation reduction potentials. Plant disease. 2012 Jun;96(6):875–80.

17 Gryzenhout, M. (2010). Mushrooms of South Africa. Struik Nature, Cape Town, South Africa.

18 Marx DH. The influence of ectotrophic mycorrhizal fungi on the resistance of pine roots to pathogenic infections. I. Antagonism of mycorrhizal fungi to root pathogenic fungi and soil bacteria. Phytopathology. 1969;59:153–63.

19 White TJ, Bruns T, Lee SJ, Taylor JL. Amplification and direct sequencing of fungal ribosomal RNA genes for phylogenetics. PCR protocols: a guide to methods and applications. 1990 Jan 1;18(1):315–22.

20 Kumar S, Stecher G, Tamura K. MEGA7: molecular evolutionary genetics analysis version 7.0 for bigger datasets. Molecular biology and evolution. 2016 Mar 22;33(7):1870–4.

21 Altschul SF, Madden TL, Schäffer AA, Zhang J, Zhang Z, Miller W, Lipman DJ. Gapped BLAST and PSI-BLAST: a new generation of protein database search programs. Nucleic acids research. 1997 Sep 1;25(17):3389–402.

22 Nilsson RH, Larsson KH, Taylor AF, Bengtsson-Palme J, Jeppesen TS, Schigel D, Kennedy P, Picard K, Glöckner FO, Tedersoo L, Saar I. The UNITE database for molecular identification of fungi: handling dark taxa and parallel taxonomic classifications. Nucleic acids research. 2018 Oct 29;47(D1):D259–64.

23 Tennant D. A test of a modified line intersect method of estimating root length. The Journal of Ecology. 1975 Nov 1:995–1001.

24 Team R. RStudio: integrated development for R. RStudio, Inc., Boston, MA URL http://www.rstudio.com. 2015 Jun;42:14.

25 Mateos E, Olaizola J, Pajares JA, Pando VI, Diez JJ. Influence of *Suillus luteus* on *Fusarium* damping-off in pine seedlings. African Journal of Biotechnology. 2017 Feb 8;16(6):268–73.

26 Mucha J, Dahm H, Strzelczyk E, Werner A. Synthesis of enzymes connected with mycoparasitism by ectomycorrhizal fungi. Archives of microbiology. 2006 Mar 1;185(1):69–77.

27 Soytong K, Sibounnavong P, Kanokmedhakul K, Kanokmedhakul S. Biological active compounds of *Scleroderma citrinum* that inhibit plant pathogenic fungi. Journal of Agricultural Technology. 2014;10(1):79–86.

28 Takakura Y. *Tricholoma matsutake* fruit bodies secrete hydrogen peroxide as a potent inhibitor of fungal growth. Canadian journal of microbiology. 2015 Mar 6;61(6):447–50.

29 Duchesne LC, Peterson RL, Ellis BE. Interaction between the ectomycorrhizal fungus *Paxillus involutus* and *Pinus resinosa* induces resistance to *Fusarium oxysporum*. Canadian Journal of Botany. 1988 Mar 1;66(3):558–62.

30 Duchesne LC, Peterson RL, Ellis BE. Pine root exudate stimulates the synthesis of antifungal compounds by the ectomycorrhizal fungus *Paxillus involutus*. New Phytologist. 1988 Apr 1;108(4):471–6.

31 Bennett AE, Alers-Garcia J, Bever JD. Three-way interactions among mutualistic mycorrhizal fungi, plants, and plant enemies: hypotheses and synthesis. The American Naturalist. 2005 Dec 19;167(2):141–52.

